# Effects of Agroforestry Trees on Microclimate and Enset (*Ensete ventricosum*) Morphophysiology in South Ethiopia

**DOI:** 10.64898/2026.03.23.713702

**Authors:** Admasu Yirgalem, Gezahegn Garo, Rony Swennen, Sabura Shara, Bart Muys, Oliver Honnay, Karen Vancampenhout

**Author notes:** Corresponding author. Division for Forest, Nature and Landscape, KU Leuven, Campus Geel, Belgium.

## Abstract

Enset (*Ensete ventricosum*), a multipurpose crop domesticated exclusively in Ethiopia, serves as a staple food for millions of smallholder farmers. It is primarily cultivated as a monocrop in homegardens, leaving it vulnerable to climate change risks. One potential nature-based solution involves agroforestry systems; however, enset’s response to canopy cover remains unclear. This study examined how scattered trees in enset farms affected microclimate and enset morpho-physiology in South Ethiopia. Trees significantly modified the microclimate conditions in enset homegardens. The average daily reductions in air, soil surface, and soil temperatures ranged from -0.5 to -1.9 °C, -0.4 to -2.1 °C, and +0.4 to -1.0 °C, respectively. The minimum soil moisture offset ranged from +0.8% to +5.7%. Although the tree identity effect on enset growth was negligible, planting position relative to the overstory trees significantly influenced enset responses. Most morphophysiological traits were higher under tree canopies, with progressively lower values at the edge and outside the tree canopy. In contrast, leaf dry matter content exhibited an inverse trend, aligning with the leaf economics spectrum. These results demonstrate enset’s adaptability to canopy shade, suggesting potential for agroforestry expansion. Cultivar-specific shade tolerance and ideal shade levels to maintain enset productivity should be investigated further.

## 1. Introduction

In 2023, an estimated 28.9% of the global population was moderately or severely food insecure (FAO *et al*., 2024). This situation has worsened since 2015, mainly due to conflicts and climate-related extremes, jeopardizing the progress toward achieving the UN Sustainable Development Goal of zero hunger by 2030 (IPCC, 2023). Especially in sub-Saharan Africa, the prevalence of severe food insecurity is on the rise (FAO *et al*., 2024), and the population of 1.2 billion now is expected to surpass 2 billion by 2050, increasing food demand by 3.9 percent per year (Cardell *et al*., 2024). It is key that increasing agricultural production happens sustainably, without undermining the ecosystems’ capacity to sustain human well-being (Torres *et al*., 2021). In this context, neglected/orphan crops are gaining attention as an avenue to alleviate food insecurity challenges under a changing climate (Talabi *et al*., 2022). Often cultivated by subsistence farmers, these crops have significant potential to contribute to sustainable food systems (Mabhaudhi *et al*., 2019; Tadele *et al*., 2024). They are often more stress-tolerant and may have unique and beneficial nutritional profiles (Chapman *et al*., 2022). Enset (*Ensete ventricosum*), a multipurpose perennial herbaceous plant domesticated only in Ethiopia, is a prime example of an underexploited crop and a potential candidate for famine insurance in climate-vulnerable regions (African Orphan Crops Consortium, 2013; FAO *et al*., 2024; National Research Council, 2006). Despite being a staple and food security crop for millions (Borrell *et al*., 2020; Kidane *et al*., 2021; Tadele *et al*., 2024), enset has been relatively neglected by scientific research and is inarguably the least-studied African crop (Venkatesan *et al*., 2022). Known as the ‘tree against hunger’ (Brandt *et al*., 1997), enset offers flexible harvest timing, high yield, long storage, and putative drought tolerance, enabling farmers to bridge adverse periods of harsh conditions (Birmeta *et al*., 2004; Borrell *et al*., 2020; Chase *et al*., 2023; Yemata, 2020). It is considered a first-rated climate-smart crop due to its ability to withstand prolonged drought lasting more than five years (Kudama *et al*., 2022). Due to this capability, areas with a more frequent severe drought history in southwestern Ethiopia have a much larger proportion of enset production (Chase *et al*., 2023). According to Tsegaye & Struik (2001), its potential yields per hectare may be higher than any other crop cultivated in Ethiopia. Enset-producing households in Ethiopia are therefore less vulnerable to shocks and perceive less risk (Feyisa *et al*., 2022). Enset is now estimated to be a staple crop feeding 20 million people in its current cultivation range in Ethiopia. But its wild distribution suggests that it could serve a much wider area in Ethiopia, Kenya, Uganda, and Rwanda, and enset cultivation might prove feasible for an additional 87.2–111.5 million people (Koch *et al*., 2021).

The enset production system in most parts of southern Ethiopia is a monoculture in homegardens (Degefa & Dawit, 2018), surrounding houses (Shara *et al*., 2021). Although enset requires relatively few off-farm inputs (Senbeta *et al*., 2022), those further from the house often remain undernourished due to limited manure application, resulting in reduced growth because of low soil nutrient availability and reduced organic matter, higher temperature and moisture stress, or due to perceptions that preferred food types need less input (Amede & Taboge, 2007; Shara *et al*., 2021). Ongoing climate change in the Central Rift of Ethiopia is likely to exacerbate this, which makes adaptation strategies increasingly urgent (Belay *et al*., 2017). The introduction of trees into farming systems (agroforestry) has the potential to address these climate change-related challenges (Torres *et al*., 2021). Indeed, the potential benefits of agroforestry are well-known. For example, compared to monocropping, agroforestry systems are better buffered against extreme climate events like temperature fluctuations (Niether *et al*., 2018). Additionally, agroforestry supports long-term productivity (IPCC, 2023) and diversifies farm income (Kassie, 2017). Agroforestry systems in East Africa contribute to livelihoods by providing food, fodder, firewood, and income (Muthuri *et al*., 2023). Moreover, these practices help to reduce runoff and soil loss and improve slope stability, infiltration rates, and soil moisture content (Hemp, 2006; Kuyah *et al*., 2019). Enset is a monocotyledonous crop with a fibrous rooting system that grows from the corm and primarily exploits nutrients from the topsoil (Brandt *et al*., 1997; Zewdie *et al*., 2008). Woody trees, on the other hand, enhance soil fertility through nutrient cycling and enrich the soil through litter deposition (Negash & Starr, 2013), which therefore enhances nutrient availability. Enset wild relatives are reported to grow in forest gaps (Birmeta *et al*., 2004; Borrell *et al*., 2019), and interestingly, in some parts of Southern Ethiopia, it is also cultivated in homegarden agroforestry systems (Abebe & Bongers, 2012; Mellisse *et al*., 2018). Those systems, which were originally dominated by natural forests, integrated *Ensete ventricosum* and *Coffea arabica* following selective canopy thinning (Negash & Starr, 2013). Thus, fields are enriched with diverse, multipurpose tree species that can provide food and agroecological services, such as improving soil fertility, controlling erosion, mitigating climate change, and conserving biological diversity (Abebe *et al*., 2006; Lelamo, 2021; Wolka *et al*., 2021). However, such traditional enset-based homegardens are in decline due to socioeconomic changes (Abebe, 2018; Sahle *et al*., 2022). Nevertheless, these traditional systems suggest that it is possible to grow enset under a (light) canopy of trees and benefit from the added benefits of agroforestry.

While the general benefits of introducing trees in cropping systems are well-known, tree-crop interactions can also lead to trade-offs (Gonçalves *et al*., 2021). Such trade-offs often arise from competition between trees and crops for space, water, nutrients, and light (Tschora & Cherubini, 2020), resulting in a reduction in the biomass harvest of the main crop (Bertsch-Hoermann *et al*., 2021; Niether *et al*., 2020). Among these factors, competition for light is recognized as a critical factor (Scordia *et al*., 2023) and it is strongly influenced by the canopy structure of the tree species, which affects air temperature, relative humidity, wind speed, and solar radiation intensity (Feng *et al*., 2023; Speak *et al*., 2020). Despite the enset’s potential importance, relatively little is known about its biology and ecology (Borrell *et al*., 2019). The wide range of suitable agroecological conditions for enset cultivation suggests a large phenotypic plasticity to changes in the environment, such as elevation (Negash, 2001; Yemataw *et al*., 2018). Such plasticity enables species to thrive in heterogeneous environments (Nürnberger, 2013; Sommer, 2020) and can facilitate subsequent adaptive evolution (Lalejini *et al*., 2021; West-Eberhard, 2008). However, despite the enset’s traditional cultivation under tree canopies in agroforestry systems, there is hardly any data on its interaction with trees, particularly regarding responses to canopy cover remains poorly documented.

Here, we capitalized on the presence of woody tree species in enset-producing homegardens in the Gamo highlands of South Ethiopia to evaluate the potential benefits of these trees in enset production systems. The specific objectives were twofold: first, to examine the effects of canopy trees of different species in enset homegardens on microclimate regulation; and second, to evaluate the effects of the tree species canopy cover on the morphophysiological traits of enset. We hypothesized that the different tree species vary in canopy structure, governing the understory enset plants’ exposure to light and the overall microclimate. Additionally, we hypothesized that enset exhibits phenotypic plasticity in response to shading imposed by the canopy cover of the tree species. Our findings supports the promotion of enset-based homegarden agroforestry systems by addressing knowledge gaps related to enset’s phenotypic interaction with woody tree species.

## 2. Material and methods

### Description of the study area

The study was conducted in Chencha Zuria, Gerese Zuria, and Kamba Zuria woredas of the Gamo highlands of south Ethiopia at an elevation ranging from 2100 to 3000 m a.s.l. The Gamo zone is geographically located between 5°55′ N and 6°20′ N latitude and between 37°10′ E and 37°40′ E longitude in the South Ethiopian Rift (Shalishe *et al*., 2023).

The mean temperature ranges between 23°C and 14°C in the lowlands and highlands, respectively; the mean annual rainfall is between 750mm and 1700mm (Berhanu *et al*., 2013; Coltorti *et al*., 2019). The landscape is characterized by steep slopes affected by landslides and dissected by concave valleys and gullies, with the uppermost part characterized by gently undulating surfaces across a range of altitudes (Coltorti *et al*., 2019). The region’s natural vegetation is Afromontane forest; however, the landscape has been severely deforested and is now predominantly covered by annual crops (Assefa & Bork, 2014, 2016). The Gamo people are predominantly farmers, with mixed crops and livestock production (Gemeda *et al*., 2023). Enset is among the main food crops and is grown in traditional homegardens surrounding the house (Shara *et al*., 2021). Unlike the typical enset-agroforestry systems in the Gedeo and Sidama Zones, in Gamo, trees are rarely incorporated into enset plots and are instead mainly found along farm boundaries, near roads, or in small patches of woodlands (own observation).

### Study species selection

A reconnaissance field visit was conducted in the Gamo highlands’ enset-producing homegardens in three woredas to identify the commonly present tree species and the availability of trees integrated with the enset farming system. During the survey, over 150 enset homegardens were randomly selected for observation, and seven tree species were identified as commonly present. These species, listed in order of importance/dominance, are *Croton macrostachyus, Ficus sur, Erythrina brucei, Hagenia abyssinica, Cordia africana, Persea americana*, and *Prunus africana*.

### Experimental setup

In each homegarden, a solitary tree was selected, and three circular plots with different radii were laid out under the tree canopy. Enset leaf sampling and data recordings were done at the middle of the tree canopy, at the edge of the tree canopy, and outside the tree canopy. The selected trees were at least 10 meters away from the nearest woody plants, and the direction towards the houses was omitted from sampling, as it is reported to be often overfertilized with manure (Shara *et al*., 2021).

### Data collection

Tree age was estimated through interviews with household heads or farmers, while other biometric attributes were measured directly. Diameter at breast height (DBH) was measured using a diameter tape or a caliper, depending on the size of the tree trunk. The crown diameter (CD) was measured using a tape, and two measurements were made perpendicularly. Then, the crown area (CA) was estimated from the CD, assuming a circular crown shape (Snowdon *et al*., 2001). The variables of interest in this study were the tree canopy gap light transmission (canopy openness/closure) and CA. Canopy openness (CO) refers to the proportion of sky unobscured by vegetation/canopy (Jennings *et al*., 1999). It was assessed by taking hemispherical photographs with an Android smartphone equipped with a fisheye lens and analysed with Gap Light Analysis Mobile App, following the methods described by Cameron *et al*. (2021) and Díaz (2022). The photographs were taken under the tree canopy, and the images were visually inspected to ensure that they were fully under the canopy. For microclimate data recording, 29 farms were sub-selected from the 38 farms for feasibility reasons. Temperature-moisture-sensor (TMS) data loggers (model: standard TMS4) were used to measure microclimate parameters at 15-minute intervals starting from February 2024 to the end of July 2024. They recorded temperature at +15, 0, and −8 cm relative to the soil surface (further referred to as air, surface, and soil temperature) and volumetric soil moisture to a depth of approximately 14 cm (Wild *et al*., 2019). They were installed both under the canopy and outside the canopy of the selected tree species. From the 38 homegarden, three enset plants of two to three years old were selected at each of the three distances (a total of 342 individual enset plants). Fluorometric parameters namely maximum quantum efficiency of Photosystem II (Fv/Fm) and performance index (PIabs), and leaf chlorophyll content were measured from the third active leaf at the lamina’s central position using a portable Handy PEA (Hansatech Instruments Ltd., King’s Lynn, Norfolk, England) and a chlorophyll content meter (model: CCM-200 plus), respectively. The Fv/Fm is a sensitive indicator of plant photosynthetic performance, with healthy samples achieving a maximum value of approximately 0.85. While PI_*abs*_ is an indicator of sample vitality, indicating an internal force of the sample to resist constraints from the outside. All PI_*abs*_measurements were multiplied by 10 as a correction factor for error in the handy PEA software (Shara *et al*. 2022, PhD dissertation). Then, the middle section of the lamina was collected from these leaves following Shara *et al*. (2021), placed in an airtight plastic bag, and moved to the AMU laboratory for further analysis of leaf moisture and dry matter content and specific leaf area (SLA). A similar procedure as Garnier *et al*. (2001) was followed with little modification for SLA. A 30 cm*15 cm (450 cm^2^) section of the laminal leaf was taken from the middle part, and fresh weight was recorded. Then, these leaves were oven-dried at 60 °C for 24 hours, and then the dry mass was weighed. We then calculated specific leaf moisture content (LMoC), dry matter content (LDMC), and specific leaf area (SLA) from the fresh mass and dry mass of the section of a leaf blade. LDMC was calculated as the ratio of the dry mass of a leaf to its fresh mass and expressed as a percentage, while SLA was measured as the leaf area per dry mass of a specific leaf section.

### Statistical analysis

All statistical analyses were conducted using the R software version 4.3.3 (R Core Team, 2024). A simple GLM was used to assess the variation between tree species in terms of their biometric characteristics. Tree age was included as a covariate factor as it highly varies between trees and tree species, which could influence the canopy characteristics. The microclimate offset values were calculated as the differences between the temperatures under and outside the tree canopy, as well as the differences in soil moisture content. First, the raw microclimate data, recorded at 15-minute intervals, were processed to obtain the daily minimum and maximum values. Then, the offsets were calculated from these values. Next, we analyzed the effects of tree species on the offsets using a GLM, by including the crown area as a covariate. Then, a linear mixed-effect model was employed to model leaf moisture and leaf dry matter content, chlorophyll content, specific leaf area, quantum efficiency, and performance index against tree species, planting position relative to the tree canopy (hereafter referred to as radial distance), and their interaction, as fixed factors by individual farms as random factor. Multiple comparisons were performed with Tukey’s post hoc test using the ‘emmeans’ function in the ‘emmeans’ R package. The visualization of the results was performed using the ‘*ggplot2*’ and ‘*ggpubr*’ functions in R software.

## 3. Results

### 3.1. Tree biometric characteristics

The tree species significantly differed in mean age (F = 3.65; p < 0.05), with *F. sur* having the oldest trees, whereas the youngest trees observed were *C. macrostachyus* (Table 2). The GLM result showed no significant difference in DBH, CD, and CA between the tree species. However, significant differences were observed in CO (F = 3.71; p < 0.05) among the species. The highest CO (18.2%) was recorded for *C. macrostachyus*, which was significantly higher than the CO of *E. brucei* (13.9%) but not different from that of *F. sur* (14.7%), *C. africana* (14.8%), and *H. abyssinica* (17.1%) (Table 2).

**Table 1.** Selected tree species planted in Enset homegardens of Gamo highlands, their family, ecological services, and characteristics (Kuria *et al*., 2017)

| Tree species | Family | Ecological Services | Growth rate |
| --- | --- | --- | --- |
| *Croton macrostachyus* | Euphorbiaceae | Live fence, shade, erosion control, riverbank stabilization, windbreak, mulching | Fairly fast-growing |
| *Ficus sur* | Moraceae | Live fence, shade, erosion control | Unknown |
| *Erythrina brucei* | Fabaceae | Ornamental, live fence, erosion control, soil fertility improvement through Nitrogen-fixing, and mulch/leaves | Slow growing |
| *Hagenia abyssinica* | Rosaceae | Ornamental, live fence, erosion control, soil fertility improvement through mulch/leaves | Slow growing |
| *Cordia africana* | Boraginaceae | Ornamental, live fence, dead fence, shade, erosion control, soil fertility improvement through mulch/leaves | Reasonably fast-growing |

**Table 2.**
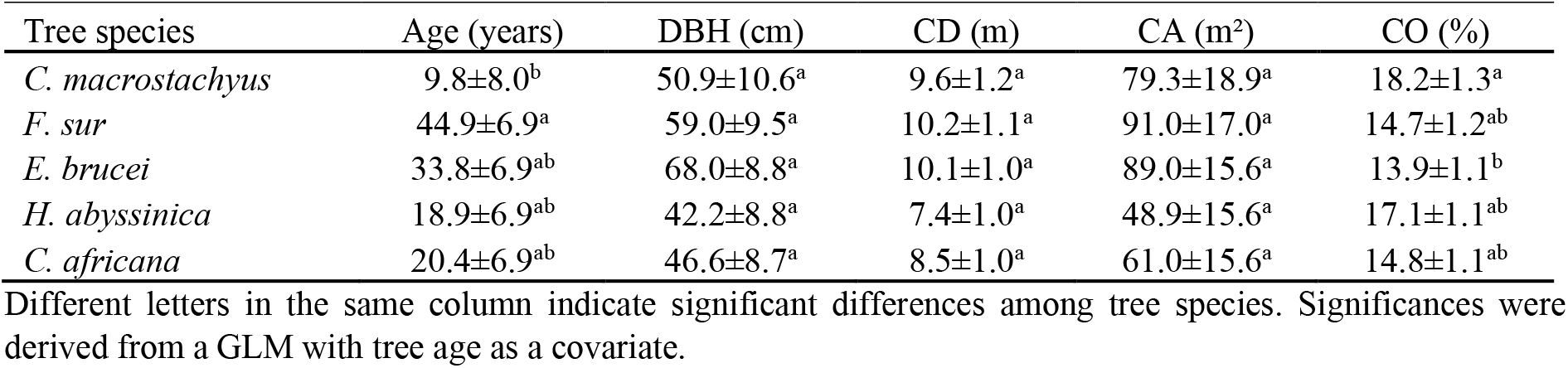
Comparison of biometric characteristics (mean±SE) of trees belonging to five species, solitarily present inside enset farms in homegardens of the Gamo highlands of South Ethiopia.

| Tree species | Age (years) | DBH (cm) | CD (m) | CA (m<sup>2</sup>) | CO (%) |
| --- | --- | --- | --- | --- | --- |
| *C. macrostachyus* | 9.8±8.0<sup>b</sup> | 50.9±10.6<sup>a</sup> | 9.6±1.2<sup>a</sup> | 79.3±18.9<sup>a</sup> | 18.2±1.3<sup>a</sup> |
| *F. sur* | 44.9±6.9<sup>a</sup> | 59.0±9.5<sup>a</sup> | 10.2±1.1<sup>a</sup> | 91.0±17.0<sup>a</sup> | 14.7±1.2<sup>ab</sup> |
| *E. brucei* | 33.8±6.9<sup>ab</sup> | 68.0±8.8<sup>a</sup> | 10.1±1.0<sup>a</sup> | 89.0±15.6<sup>a</sup> | 13.9±1.1<sup>b</sup> |
| *H. abyssinica* | 18.9±6.9<sup>ab</sup> | 42.2±8.8<sup>a</sup> | 7.4±1.0<sup>a</sup> | 48.9±15.6<sup>a</sup> | 17.1±1.1<sup>ab</sup> |
| *C. africana* | 20.4±6.9<sup>ab</sup> | 46.6±8.7<sup>a</sup> | 8.5±1.0<sup>a</sup> | 61.0±15.6<sup>a</sup> | 14.8±1.1<sup>ab</sup> |

Different letters in the same column indicate significant differences among tree species. Significances were derived from a GLM with tree age as a covariate.

### 3.2. Tree influences on microclimate offsets in enset homegardens

The daily offsets of maximum air, surface, and soil temperatures were highly significantly influenced by tree species (Table 3). The highest maximum air temperature offset was recorded under *F. sur* (-1.9 °C), while the lowest offset was recorded under *H. abyssinica* (-0.5 °C) (Figure 3, Table S1). Similarly, the highest and lowest cooling of the maximum surface temperature was recorded under *F. sur* (-2.1 °C) and *H. abyssinica* (-0.4 °C), respectively. On the other hand, the highest maximum soil temperature offset was recorded under *F. sur* (-1.0 °C), which was not significantly different from the offset by *C. africana* (-0.9 °C) and *C. macrostachyus* (-0.8 °C). The lowest offset (+0.4 °C) was recorded under *H. abyssinica* (Figure 3, Table S1). Tree species significantly influenced the minimum volumetric soil moisture content (VMC) offsets (p < 0.001, Table 3). The highest soil moisture offset was observed under *E. brucei* (+5.7 %), while the lowest minimum soil moisture offset (+0.8%) was recorded under *H. abyssinica* (Figure 3, Table S1). On the other hand, the soil moisture offsets observed by *C. africana, C. macrostachyus*, and *F. sur* were statistically similar (Figure 3).

**Figure 1.**
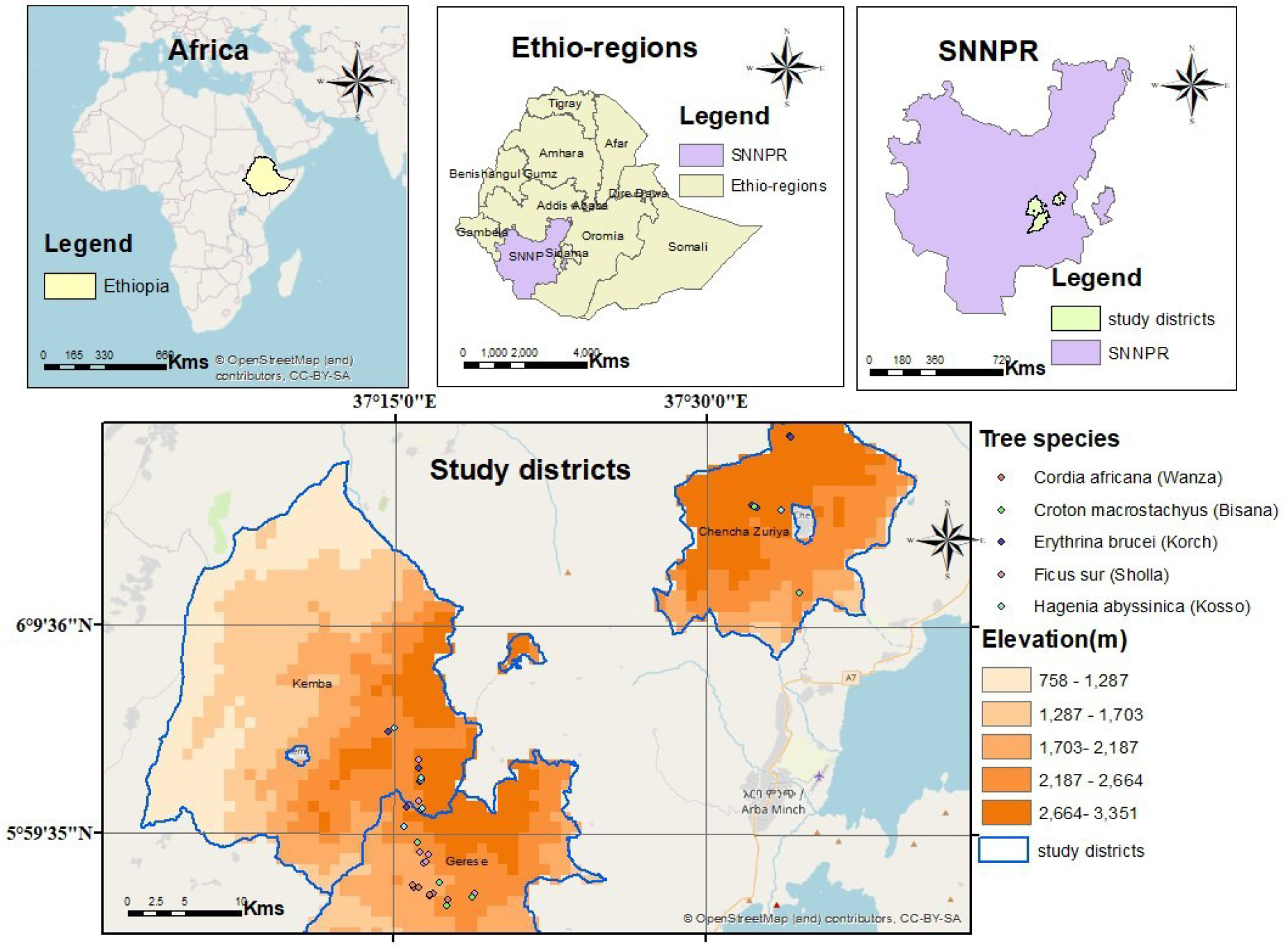
Sampling plots (individual enset farms) in three woredas (Chencha Zuria, Gerese Zuria, and Kemba Zuria) of the Gamo highlands of South Ethiopia.

**Figure 2.**
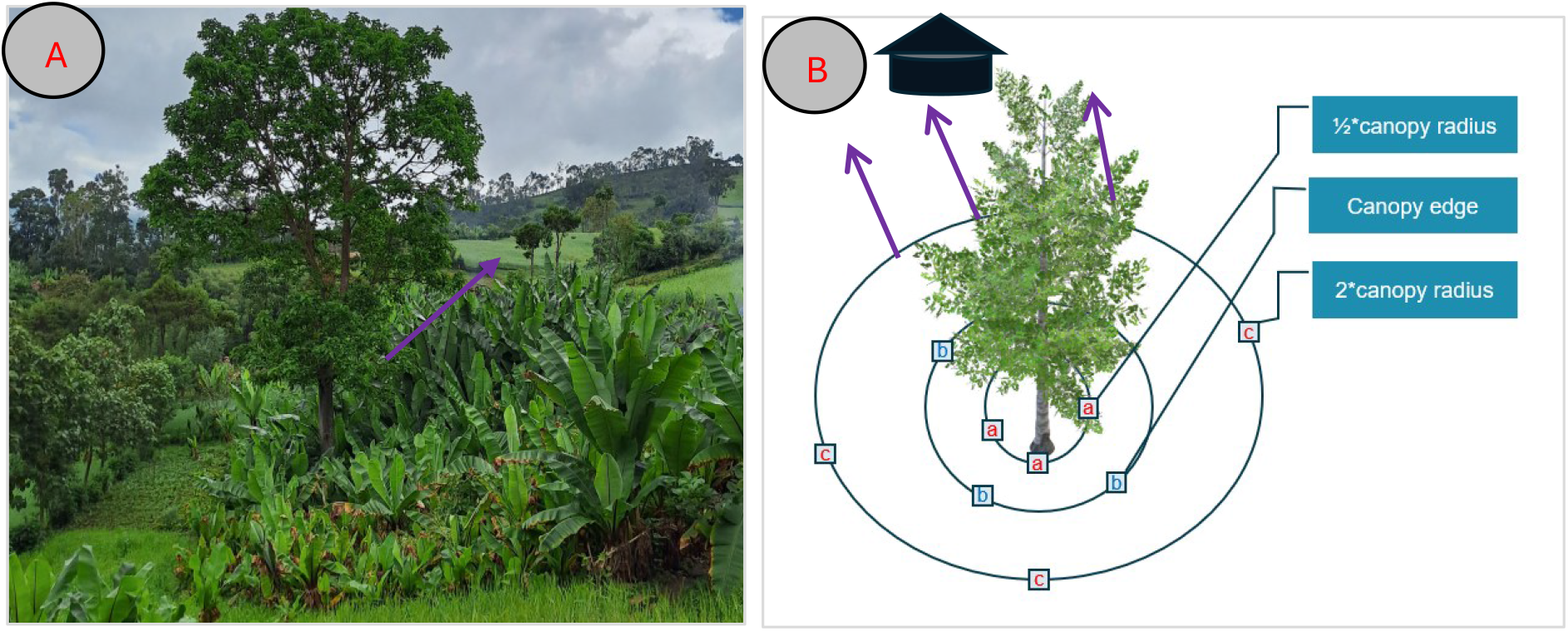
Enset farm with a solitary tree (A) and sampling scheme along three distances from a tree trunk (B). Enset data recording and sampling were done in the middle of the canopy (a), at the edge (b), and outside the canopy (c). Microclimate data recording loggers were installed in the middle and outside of the tree canopy for six months (February 2024 to July 2024). Arrows indicate direction towards the houses omitted from sampling.

**Figure 3.**
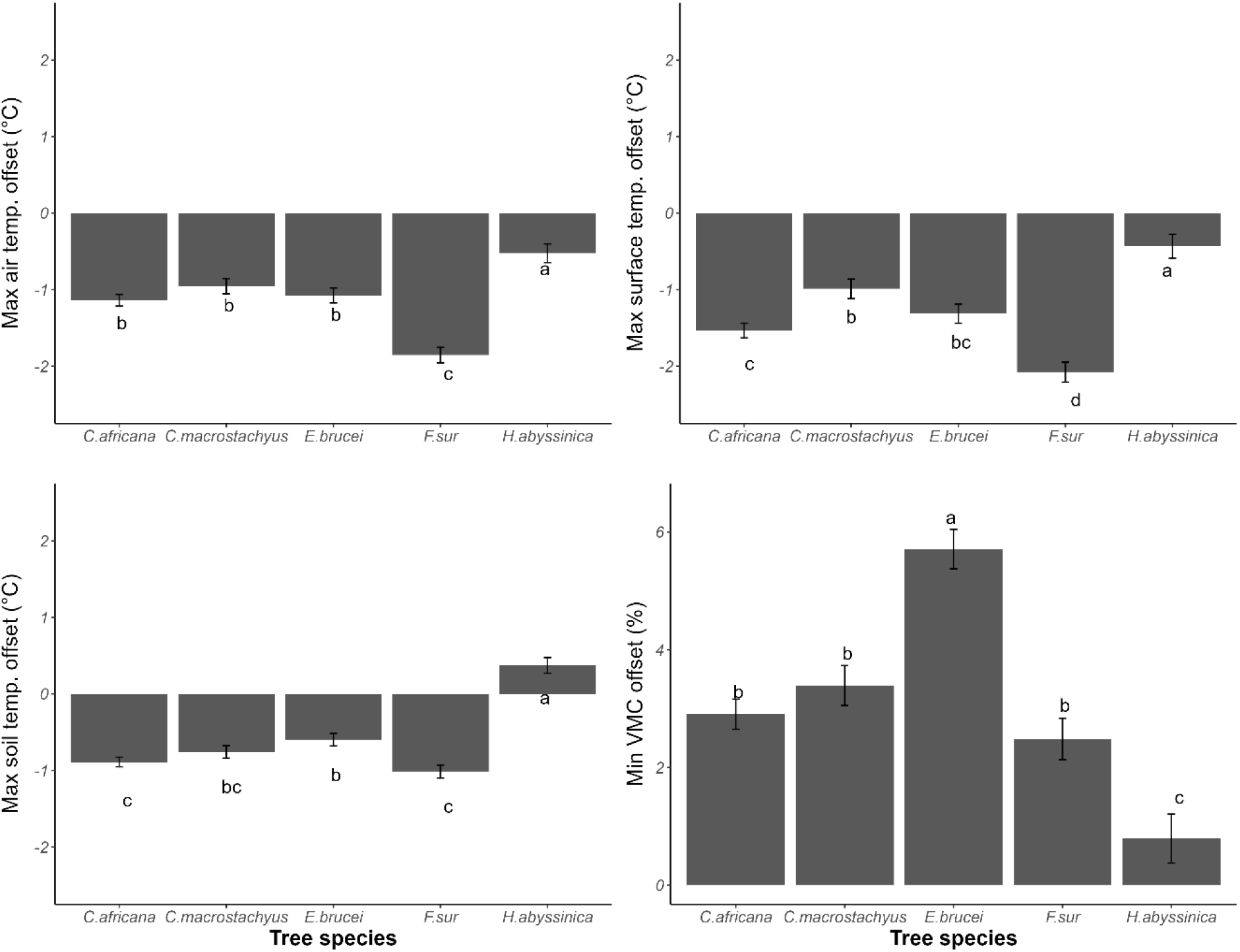
Microclimate offsets under solitary trees of five tree species in enset homegardens of the Gamo highlands of South Ethiopia. Bars represent the offsets of maximum air, surface, and soil temperature, and soil volumetric moisture content offsets. Different letters in each figure indicate significant differences among the tree species’ effects on offsets based on a Tukey’s HSD post hoc test. VMC: soil volumetric moisture content.

**Table 3.**
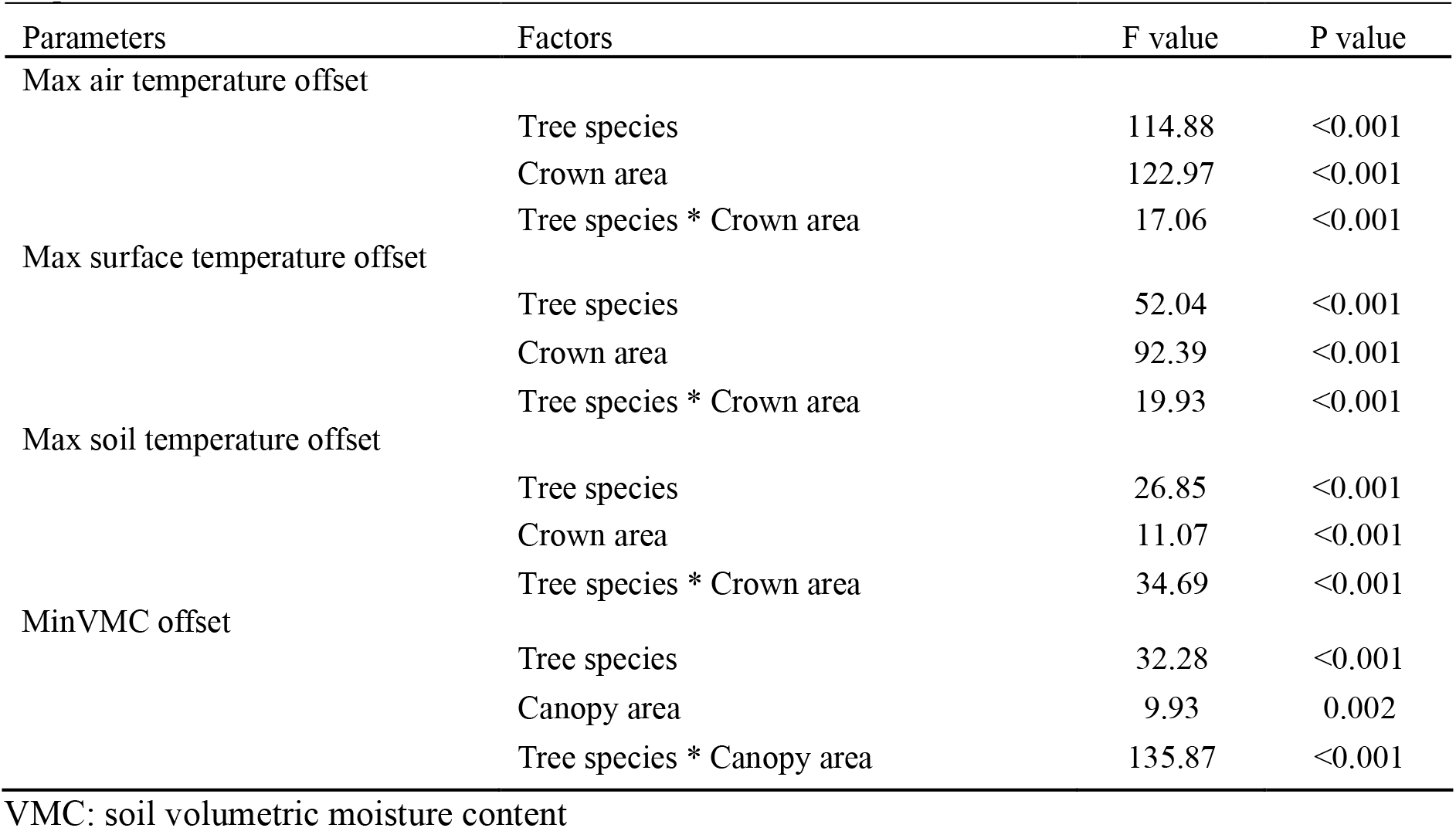
The effect of tree species on offsets of microclimate parameters (maximum air, surface, and soil temperature, and soil moisture content) with the tree’s crown area is considered a covariate.

All the microclimate offsets were also significantly affected by the interaction between tree species and crown area (p < 0.001) (Table 3), indicating that the positive effects of increasing crown area on offsets were highly dependent on the tree species (Fig S1). An increase in crown area increased the temperature offsets, with *C. africana* providing the strongest temperature offsets. Exceptionally, an inverse correlation was observed for soil temperature offset by *H. abyssinica*. Regarding the soil volumetric moisture content, the offsets by *C. africana, C. macrostachyus*, and *H. abyssinica* were positively correlated with crown area, whereas negative offsets were recorded from *E. brucei* and *F. sur*.

### 3.3. Tree canopy cover effect on enset morpho-physiological traits

Leaf moisture content of enset varied significantly by tree species (F = 2.50, p = 0.06) and radial distance (F = 9.13, p < 0.001), but not by their interaction (F = 1.37, p = 0.21) (Table 4). Enset under the tree canopy had the highest moisture content (85.5%), significantly higher than outside the canopy (84.5%), but not different from the canopy edge (85.3%) (Table S2). Despite the non-significant interaction, pairwise comparisons showed that tree species affected leaf moisture in the middle of the canopy. Similarly, the interaction of tree species and radial distance had no significant effect on leaf dry matter content (F = 1.37, p = 0.21), but radial distance alone did (F = 9.13, p < 0.001) (Table 4). The highest dry matter content (15.5%) was recorded from enset outside the canopy, significantly higher than under the canopy (14.5%) (Table S2). Tree species significantly affected dry matter content only in the middle of the canopy, where it was lower under *E. brucei* than *C. africana* (Figure 5).

**Table 4.** Effects of tree species, radial distance (middle, edge, and outside canopy), and their interaction on LMoC, LDMC, SLA, leaf chlorophyll content (CCI), (Fv/Fm and PI_*abs*_ inferred by linear mixed model with farm included as a random factor.

| Response variables | F value | DF | P value | R<sup>2</sup>m/R<sup>2</sup>c |
| --- | --- | --- | --- | --- |
| LMoC |  |  |  |  |
| Tree species | 2.50 | 4 | 0.06 | 13.35/37.65 |
| Radial distance | 9.13 | 2 | <0.001 |  |
| Tree species x Radial distance | 1.37 | 8 | 0.21 |  |
| LDMC |  |  |  |  |
| Tree species | 2.50 | 4 | 0.06 |  |
| Radial distance | 9.13 | 2 | <0.001 | 13.35/37.65 |
| Tree species x Radial distance | 1.37 | 8 | 0.21 |  |
| SLA |  |  |  |  |
| Tree species | 1.62 | 4 | 0.19 |  |
| Radial distance | 21.68 | 2 | <0.001 | 15.28/39.32 |
| Tree species x Radial distance | 1.94 | 8 | 0.05 |  |
| CCI |  |  |  |  |
| Tree species | 2.09 | 4 | 0.11 |  |
| Radial distance | 2.70 | 2 | 0.06 | 13.14/30.87 |
| Tree species x Radial distance | 1.63 | 8 | 0.12 |  |
| Fv/Fm |  |  |  |  |
| Tree species | 1.80 | 4 | 0.15 |  |
| Radial distance | 14.76 | 2 | <0.001 | 12.08/22.18 |
| Tree species x Radial distance | 1.24 | 8 | 0.28 |  |
| PI<sub>abs</sub> |  |  |  |  |
| Tree species | 2.79 | 4 | 0.04 |  |
| Radial distance | 15.74 | 2 | <0.001 | 13.98/31.33 |
| Tree species x Radial distance | 0.25 | 8 | 0.98 |  |

**Figure 5.**
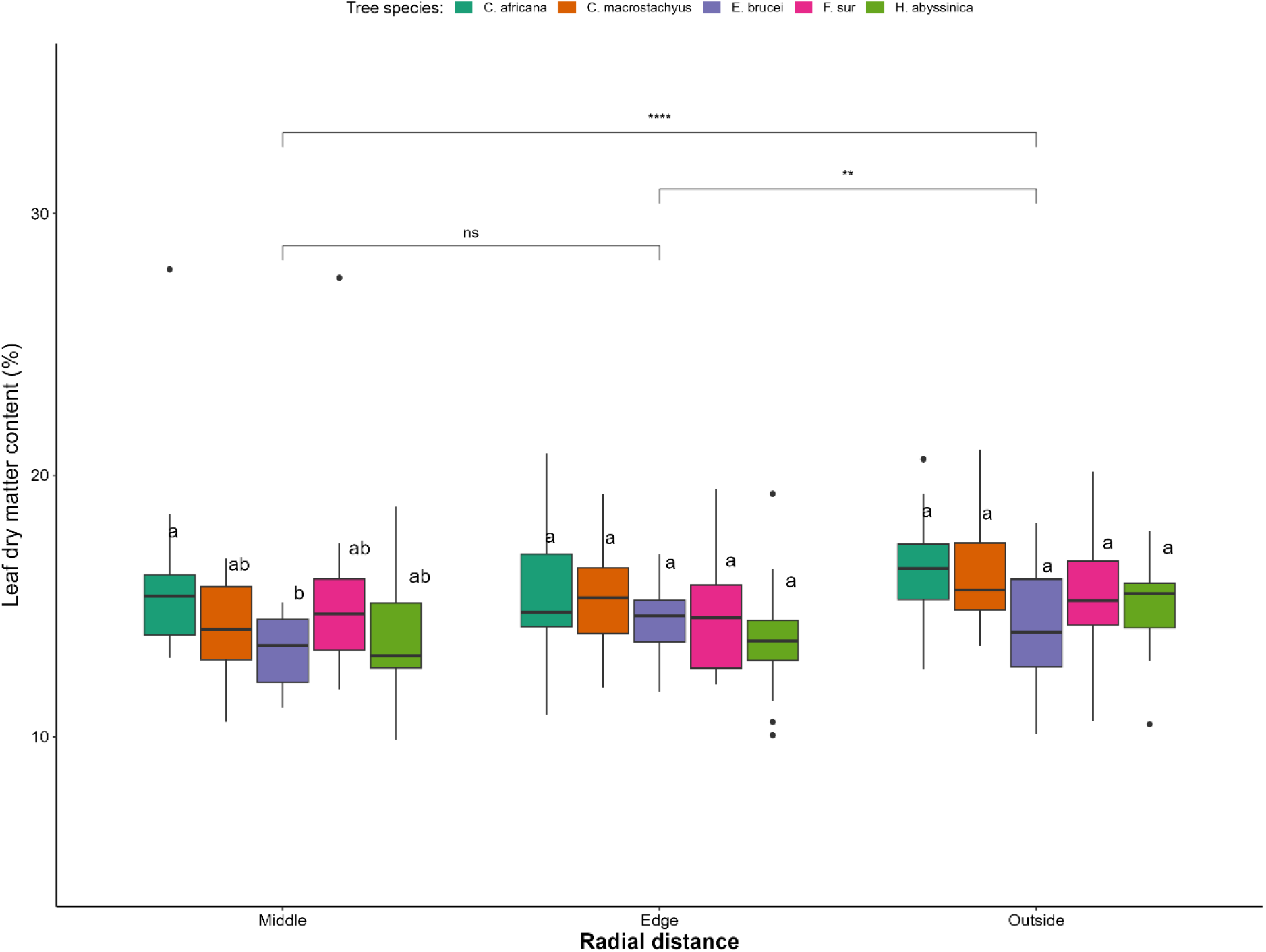
Box plots comparing tree species and enset radial distance (at the middle, edge, and outside canopy) effects on LDMC of enset grown in homegardens of Gamo highlands of Ethiopia. Different letters within each distance indicate significant differences among tree species based on a Tukey HSD post hoc test. The differences between planting positions are indicated by ‘ns’: p-value>0.05, **: p-value<0.01, ***: p-value <0.001.

Radial distance significantly affected the specific leaf area (SLA) of enset (F = 21.68, p < 0.001; Table 4). Enset outside the canopy had a lower SLA (200 cm^2^/g) than those at the edge (220 cm^2^/g) and under the canopy (229 cm^2^/g) (Table 4, Figure 6A), showing a decreasing trend with distance from the tree. However, SLA did not differ significantly between enset at the middle and the edge of the canopy. Tree species had no significant effect on SLA at any distance, except at the canopy edge, where enset at the edge of *E. brucei* had significantly lower SLA than those at the edge of *C. africana, F. sur*, and *H. abyssinica* (Figure 6A).

**Figure 6.**
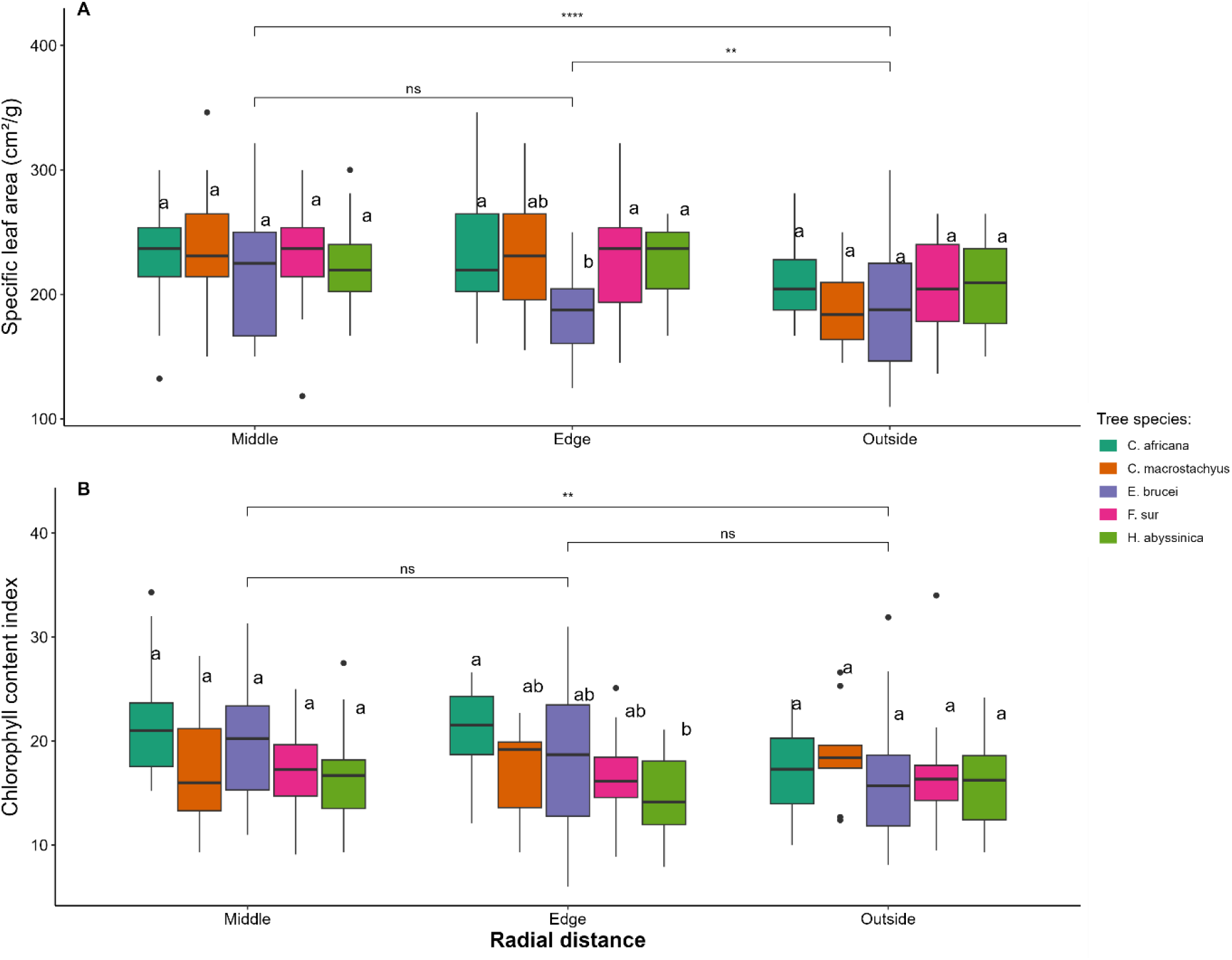
Box plots comparing tree species and radial distance (at the middle, edge, and outside canopy) effects on SLA (A) and leaf chlorophyll content index (B) of enset grown in homegardens of Gamo highlands of Ethiopia. Different letters within each distance indicate significant differences among tree species based on a Tukey HSD post hoc test. The differences between planting positions are indicated by ‘ns’: p-value>0.05, **: p-value<0.01, ***: p-value <0.001.

Leaf chlorophyll content was not significantly influenced by the interaction of tree species and radial distance (F = 2.09, p = 0.12) or by tree species alone (F = 1.63, p = 0.11) (Table 4). It was marginally affected by radial distance (F = 2.7, p = 0.06, Table 4). This marginal difference showed a slight gradient along the radial distance, with the highest chlorophyll content index (18.7) in the middle of the tree canopy, followed by the edge (17.5) and outside (17.0) (Table S2). The pairwise comparison revealed a significant effect of tree identity only at the edge of the canopy, where enset at the edge of H. abyssinica had lower chlorophyll content than at the edge of C. africana (Figure 6B).

Enset leaf chlorophyll fluorometric variables varied significantly across planting positions relative to the tree canopy (Table 4). The maximum quantum efficiency was significantly affected by the radial distance (F = 14.76, p < 0.001, Table 4), but not by tree species alone (F = 1.80, p = 0.15, Table 4) or their interaction (F = 1.24, p = 0.28, Table 4). The highest quantum efficiency (0.82) was recorded under the tree canopy, significantly higher than outside the tree canopy (0.79) (Table S2; Figure 7A). Similarly, the Fv/Fm outside the tree canopy was also significantly lower than at the edge of the tree canopy (Figure 7A). Within the radial distances, tree species had no significant effect on Fv/Fm except a marginal difference outside the canopy, where Fv/Fm outside *C. macrostachyus* was significantly lower than outside *C. africana* (Figure 7A).

**Figure 7.**
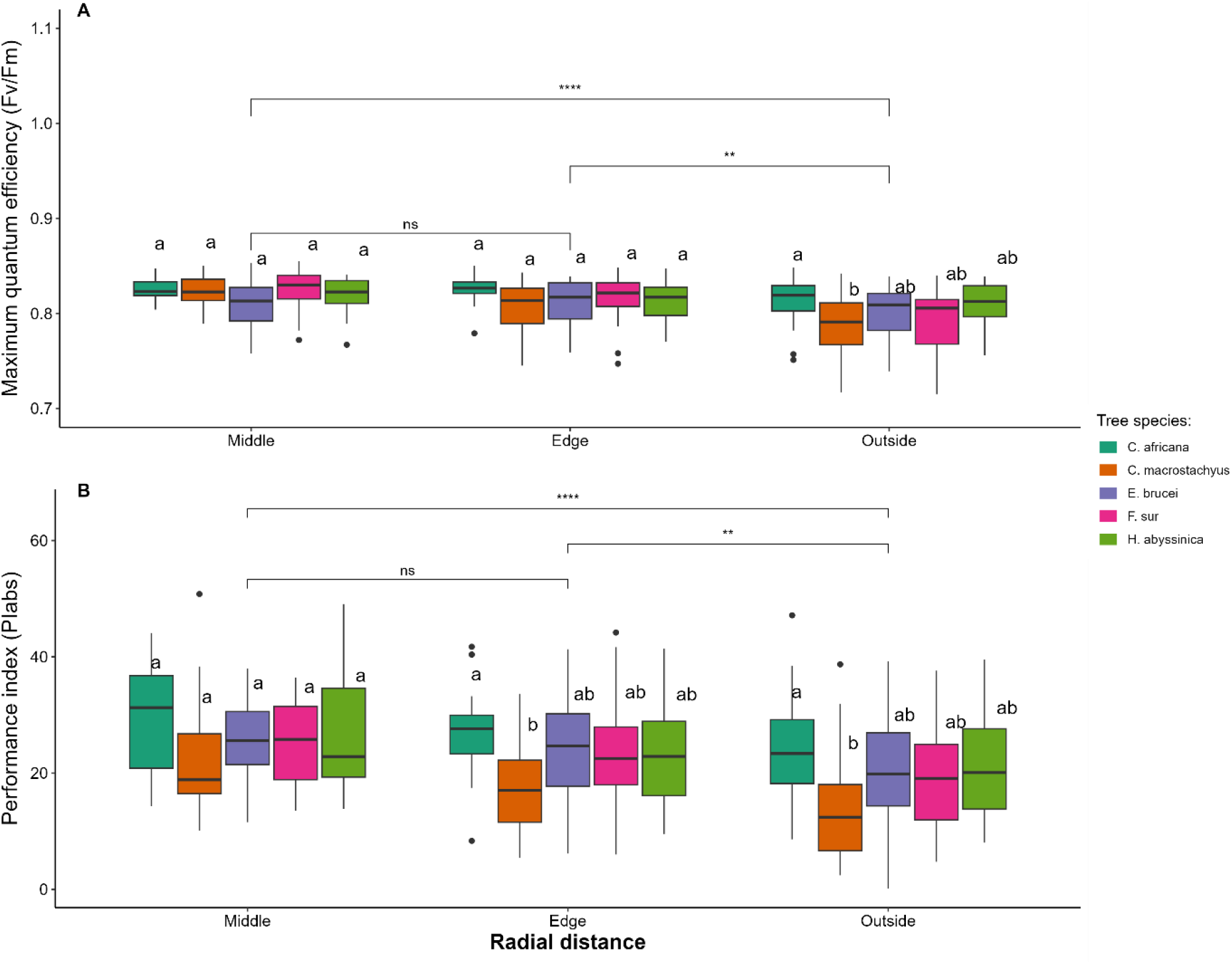
Box plots representing tree species and radial distance (at the middle, edge, and outside canopy) effects on the Fv/Fm (A) and PI_*abs*_(B) of enset grown in homegardens of Gamo highlands of Ethiopia. Different letters within each distance indicate significant differences among tree species based on a Tukey HSD post-hoc test. The differences between planting positions are indicated by ‘ns’: p-value > 0.05, **: p-value<0.01, ***: p-value <0.001.

Similarly, the performance index of enset was not significantly affected by the interaction of tree species and radial distance (F = 0.25, p = 0.98) but was significantly influenced by the tree species (F = 2.79, p < 0.05) and radial distance (F = 15.74, p < 0.001) (Table 4). The highest index (26.0) was recorded under the tree canopy, significantly higher than at the edge (23.4) and outside the tree canopy (20.0) (Table S2, Figure 7B). Generally, both Fv/Fm and PI_abs_ showed a decreasing trend with increasing distance from the tree (Table S2, Figure 8). Considering the tree species’ effect in each radial distance, a significant difference was observed at the edge and outside the canopy, with significantly lower performance indexes of enset planted at the edge and outside the canopy of *C. macrostachyus* compared to *C. africana* (Figure 7B).

## 4. Discussion

### 4.1. Tree species impact on microclimate offsets

We found that woody trees are only sparingly incorporated into the enset-based farming systems of the Gamo highlands. The common tree species identified include *C. macrostachyus, F. sur, E. brucei, H. abyssinica*, and *C. africana*, either within enset farms or in the outfields (outside the enset farms). These trees have been identified as vital native species in the agroforestry systems of Ethiopia before (Lelamo, 2021; Molla *et al*., 2023). Some have also been acknowledged as the most common tree species in banana agroforestry systems in central Uganda (Mpiira *et al*., 2013). The trees in the study area exhibited significant variation in age and canopy structure, particularly in canopy openness, which may influence understory enset performance. Specifically, the mean canopy openness followed the order: *C. macrostachyus > H. abyssinica > C. africana > F. sur > E. brucei*. This aligns with prior findings; for example, Lemenih *et al*. (2004) reported variability in canopy openness among plantation species (*Cordia africana, Eucalyptus saligna, Cupressus lusitanica*, and *Pinus patula*) in southern Ethiopia. Likewise, Kohl *et al*. (2024) found that overstory tree species significantly influenced light transmission in agroforestry systems in Ghana.

Our findings indicate that tree identity plays a key role in regulating the microclimate, closely linked to canopy characteristics such as openness and crown area. All studied species reduced both air temperature at 15cm (−0.5 °C to −1.9 °C) and surface temperature (−0.4 °C to −2.1 °C) beneath their canopies. While soil temperature reductions at 8cm depth were modest (−0.6 °C to −1.0 °C), more pronounced effects may occur under denser canopies. Notably, soil moisture at 14 cm depth was consistently higher beneath trees (+0.8% to +5.7%), highlighting their role in enhancing topsoil moisture retention. The tree buffering capacities could be beyond the recorded values, as all measurements were taken under the enset plants’ canopy, which itself ameliorates microclimate through shading with their broad leaves (Jilo *et al*., 2021; Senbeta *et al*., 2022).

Microclimate regulation varied significantly among tree species. *F. sur* exhibited the strongest buffering capacity, achieving the highest reductions in maximum daily air and soil surface temperatures. *C. africana* and *C. macrostachyus* showed similar effectiveness in moderating soil surface temperatures. In contrast, *H. abyssinica* demonstrated the weakest temperature regulation across all measured variables (air, surface, and soil temperatures). On the other hand, the highest offset of low soil moisture content was observed under *E. brucei*, while the lowest buffering effect was recorded under *H. abyssinica*. Trees’ structural attributes play a key role in influencing microclimate buffers (Zhang *et al*., 2022), with higher basal area and canopy cover improving buffering capacity (Zellweger *et al*., 2020; Zhang *et al*., 2022). Similarly, Sharmin *et al*. (2023) reported that trees with higher leaf area index and broader canopies provided the greatest cooling benefits in Australia. In an agroforestry system, Kohl *et al*. (2024) also observed substantial variation in microclimate buffering capacity among common shade tree species in Ghana. This capacity is often reflected through reducing the daily maximum temperature and increasing the daily minimum temperature (Merle *et al*., 2022). The tree height-to-crown-base also affects the overall understory environment (Blaser-Hart *et al*., 2021).

The influence of trees on microclimate regulation in our study was strongly correlated with crown area, though species effects varied across different microclimate variables. Increasing the crown area of all tree species consistently reduced the air and surface temperatures. However, the effects on soil temperature and volumetric moisture content were less distinct. For instance, soil temperature offset was strongly correlated with the crown area expansion in *C. africana, F. sur*, and *E. brucei*, but unclear for *C. macrostachyus*. In contrast, *H. abyssinica* exhibited an inverse relationship with soil temperature offset, likely because soil temperature is affected not only by canopy cover but also by the amount of leaf litter deposited on the ground (Hou *et al*., 2020). These canopy-driven microclimate regulations can enhance broader ecosystem functioning. For instance, banana-based agroforestry systems support termite survival (Godfrey *et al*., 2017), key decomposers in tropical ecosystems, thereby improving soil fertility through accelerated litter decomposition (Anbessa & Utaile, 2024; Seidelmann *et al*., 2016). In coffee agroforestry systems, shade canopies buffer temperature extremes and humidity fluctuations, enhancing pollinator diversity (Jha *et al*., 2014). Similarly, changing environmental conditions, such as soil temperature and moisture, may also alter fungal symbionts critical for nutrient cycling (Kivlin *et al*., 2011; Tedersoo *et al*., 2020).

### 4.2. Enset functional trait responses to tree canopy cover/shade

A critical factor in evaluating a crop’s suitability for agroforestry systems is its ability to thrive under a (dappled) shade. Our findings revealed that tree species identity had negligible effects on most enset leaf traits, whereas radial distance from the trunk exerted a stronger influence. All measured traits varied significantly with radial distances, indicating that microclimate gradients, rather than tree species identity, are the primary drivers of enset’s phenotypic responses. Similar results were reported for other crops, such as banana (Senevirathna *et al*., 2008), cocoa (Isaac *et al*., 2007; Kohl *et al*., 2024), sorghum (Kessler, 1994), and wheat (Yang *et al*., 2019).

Most enset leaf traits exhibited a decreasing trend with increasing distance from the tree trunk, except for leaf dry matter content, which showed an inverse relationship. Notably, the specific leaf area (SLA) of enset was significantly higher under the tree canopies than in the open area, suggesting a shade-acclimation response, a common adaptive strategy in plants growing under low-light environments (Poorter *et al*., 2009). This morphological adjustment, characterized by thinner, broader leaves, enhances light capture efficiency through increased photosynthetic surface area per unit leaf mass (Valladares & Niinemets, 2008). The elevated SLA in shaded conditions is typically associated with improved relative growth rates (Hunt & Cornelissen, 1997; Villar *et al*., 2005), indicating that enset may optimize carbon assimilation under shade. While no prior studies have explicitly examined SLA in enset, our results align with observations in other crops, such as banana (Muhidin *et al*., 2021; Senevirathna *et al*., 2008) and cacao (Isaac *et al*., 2007). These findings support the well-documented association between high SLA and shade tolerance across plant species (Janse-Ten Klooster *et al*., 2007; Sánchez-Gómez *et al*., 2006; Liu *et al*., 2016), typically involving reduced leaf thickness and stomatal density (Israeli *et al*., 1995).

This morphological adaptation was accompanied by distinct physiological changes in leaf tissue composition. We observed higher leaf moisture content and lower LDMC under shaded conditions compared to open areas, consistent with reports that high-SLA species tend to maintain greater leaf water content (Vendramini *et al*., 2002). The increased hydration under tree canopies likely reflects reduced evaporative demand due to moderated microclimate conditions (Valladares *et al*., 2008). Conversely, the lower dry matter content suggests a strategic trade-off, where shaded leaves prioritize light interception over investment in structural compounds like lignin and cellulose (Wright *et al*., 2004). While thinner leaves enhance light capture, they may increase susceptibility to physical damage (He *et al*., 2019), a critical consideration for optimization of agroforestry systems.

Interestingly, chlorophyll content showed only marginal variation across radial distance, with a slight decreasing trend with increasing distance. This stability parallels observations in Musa spp. (Thomas & Turner, 2001), suggesting some species maintain consistent chlorophyll levels despite light variations. However, while absolute chlorophyll content remained stable, we cannot rule out potential adjustments in chlorophyll a/b ratios - a common adaptation observed in *Musa spp*. and *Hevea brasiliensis* (Senevirathna *et al*., 2003, 2008) that optimizes the photosynthetic apparatus for shade conditions.

Fluorometric measurements provided further insight into enset’s photochemical adaptations. Both maximum quantum efficiency (*Fv/Fm*) and Performance Index (PI_*abs*_) were significantly higher under tree canopies, with values declining toward open areas. These results align with findings in the banana (Senevirathna *et al*., 2008; Thomas & Turner, 2001), where direct sunlight caused greater photochemical damage than diffuse light. The elevated Fv/Fm and PI_*abs*_ under shade suggest that lower irradiance mitigates light stress while maintaining sufficient energy capture (Baker, 2008). Notably, the higher photochemical efficiency alongside the stable chlorophyll content implies that enset prioritizes photosynthetic quality (efficiency) over quantity (light absorption) under shade conditions. This strategy appears effective for early growth establishment, as reduced photoinhibition in shade promotes seedling performance (Senevirathna *et al*., 2003).

Furthermore, our findings support the idea that moderate shade in agroforestry systems does not necessarily compromise productivity but may instead induce beneficial physiological adjustments. Similar patterns have been reported in other perennial crops, where shaded environments enhance water-use efficiency and reduce heat stress (Lin, 2007). In addition, other research suggests that moderate densities of tall trees in agroforestry systems can be compatible with higher productivity of banana and cacao (Salazar-Díaz & Tixier, 2019). Similarly, enhanced plant height, leaf length, leaf number, and leaf width of banana were observed under natural shading (Muhidin *et al*., 2021). Additionally, low-density shade trees improved nutrient uptake and biomass in cocoa (Isaac *et al*., 2007). Therefore, given the enset’s long-standing adaptation to agroforestry systems, its plasticity in leaf morphology likely contributes to sustained productivity under variable light conditions, a key trait for climate-resilient cropping systems. In light of alarming global climate change, efforts of reforestation in the region need to prioritize enset-based homegarden agroforestry as a resilient, sustainable production system.

## 5. Conclusion

The scattered tree species in the enset homegardens of the Gamo highlands regulated extreme climate conditions, reducing maximum air, soil surface, and soil temperatures, while keeping minimum soil moisture content higher under the tree canopy. Notably, this buffering capacity may be higher than the recorded values compared to open areas outside the enset plot. Tree species identity had a negligible effect on enset phenology, but radial distance significantly influenced responses. All morphophysiological parameters (leaf moisture content, SLA, chlorophyll content, Fv/Fm, and performance index) increased closer to the canopy, though leaf dry matter content was significantly lower under canopies than at edges or outside. Thus, we conclude that enset’s phenotypic plasticity to adapt to environmental changes suggests it may be a shade-loving crop and, more specifically, acclimate to mild shade for optimal growth.

Given the threats of global climate change and regional vulnerability to land degradation, adopting enset-based agroforestry with strategic tree spacing to enhance enset physiological performance provides smallholders a resilient and sustainable adaptation strategy. Future work should assess tree-mediated soil fertility improvements, cultivar-specific shade tolerance thresholds, yield estimation, and nutritional composition of enset products. Moreover, further research on quantifying carbon sequestration in woody tree-enset agroforestry systems may provide insight for future scaling of enset cultivation towards outfields from its current confinement to homegardens.

## Acknowledgments

We thank the Flemish Inter-University Council for Development Collaboration (VLIR-UOS), Belgium, for funding research activities through the VLIR-IUC (inter-university cooperation) program with Arba Minch University, Ethiopia. We also want to acknowledge the support of IUC coordinators Dr. Fassil Eshetu and Prof. Roel Merckx; P5 project leader Prof. Yisehak Kechero; and AMU-IUC program team members who were instrumental in facilitating logistics. We want to extend our gratitude to the farmers for sharing information and allowing access to their farms for the study, as well as Sintayehu Tomas, Legesse Bode, Beyene Kushe, and Zelalem Aniley for their assistance in gathering field data and conducting laboratory analysis. Finally, ChatGPT was instrumental in language improvement and coding assistance.

## Supplementary materials

**Table S1.** Offsets of the air temperature at 15cm (Max AiT), the soil surface temperature (Max SuT), soil temperature at a depth of -8cm (Max SoT), and soil volumetric moisture content (Min VMC) as influenced by five woody tree species sparsely planted in enset homegardens of Gamo highland of Ethiopia.

| Microclimate offsets | Tree species |  |  |  |  |
| --- | --- | --- | --- | --- | --- |
|  | *C. macrostachyus* | *E. brucei* | *F. sur* | *H. abyssinica* | *C. africana* |
| Max AiT offset (oC) | -0.9±0.1<sup>b</sup> | -1.1±0.1<sup>b</sup> | -1.9±0.1<sup>c</sup> | -0.5±0.1<sup>a</sup> | -1.1±0.1<sup>b</sup> |
| Max SuT offset (oC) | -1.0±0.1<sup>b</sup> | -1.3±0.1<sup>bc</sup> | -2.1±0.1<sup>d</sup> | -0.4±0.1<sup>a</sup> | -1.5±0.1<sup>c</sup> |
| Max SoT offset (oC) | -0.8±0.1<sup>bc</sup> | -0.6±0.1<sup>b</sup> | -1.0±0.8<sup>c</sup> | 0.4±0.1<sup>a</sup> | -0.9±0.1<sup>c</sup> |
| MinVSM offset (%) | 3.4±0.3<sup>b</sup> | 5.7±0.3<sup>a</sup> | 2.5±0.3<sup>b</sup> | 0.8±0.4<sup>c</sup> | 2.9±0.2<sup>b</sup> |

All values are given as mean±SE, and different letters in the same row indicate significant mean differences among tree species.

**Table S2.** Enset leaf morphophysiological features as influenced by tree canopy coverage in enset homegardens of Gamo highlands of Ethiopia.

| Variable | Canopy position (radial distance) |  |  |
| --- | --- | --- | --- |
|  | Middle | Edge | Outside |
| Leaf moisture content (%) | 85.5±0.25<sup>a</sup> | 85.3±0.25<sup>ab</sup> | 84.5±0.25<sup>b</sup> |
| Leaf dry matter content (%) | 14.5±0.47<sup>a</sup> | 14.7±0.25<sup>ab</sup> | 15.5±0.25<sup>b</sup> |
| Specific leaf area (cm<sup>2</sup>/g) | 229±4.8<sup>a</sup> | 220±4.8<sup>a</sup> | 200±4.8<sup>b</sup> |
| Leaf chlorophyll content (CCI) | 18.7±0.69<sup>a</sup> | 17.5±0.69<sup>ab</sup> | 17.0±0.69<sup>b</sup> |
| Fv/Fm | 0.82±0.004<sup>a</sup> | 0.81±0.004<sup>a</sup> | 0.79±0.004<sup>b</sup> |
| PI<sub>abs</sub> | 26.0±1.01<sup>a</sup> | 23.4±1.01<sup>b</sup> | 20.0±1.01<sup>c</sup> |

All values are given as mean±SE, and different letters in the same row indicate significant mean differences between planting positions relative to trees (radial distance). Fv/Fm: maximum quantum efficiency of photosystem II; Pi: performance index

**Fig S1.**
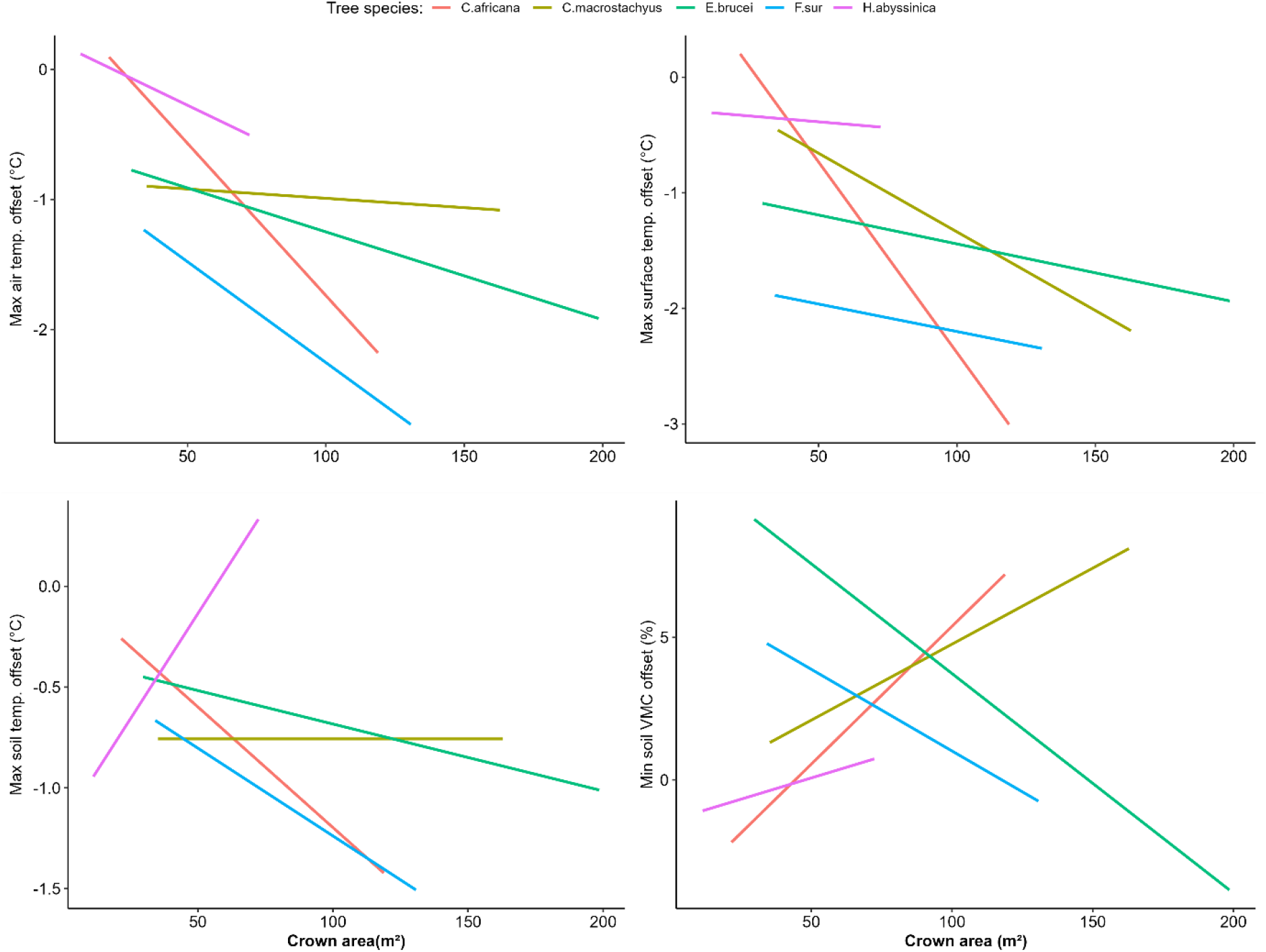
Offsets of extreme air, surface, and soil temperatures, as well as volumetric soil moisture content, as influenced by a change in the crown area for five tree species sparsely planted in enset plots of homegardens of the Gamo highlands of Ethiopia. VMC: volumetric moisture content

**Fig S2.**
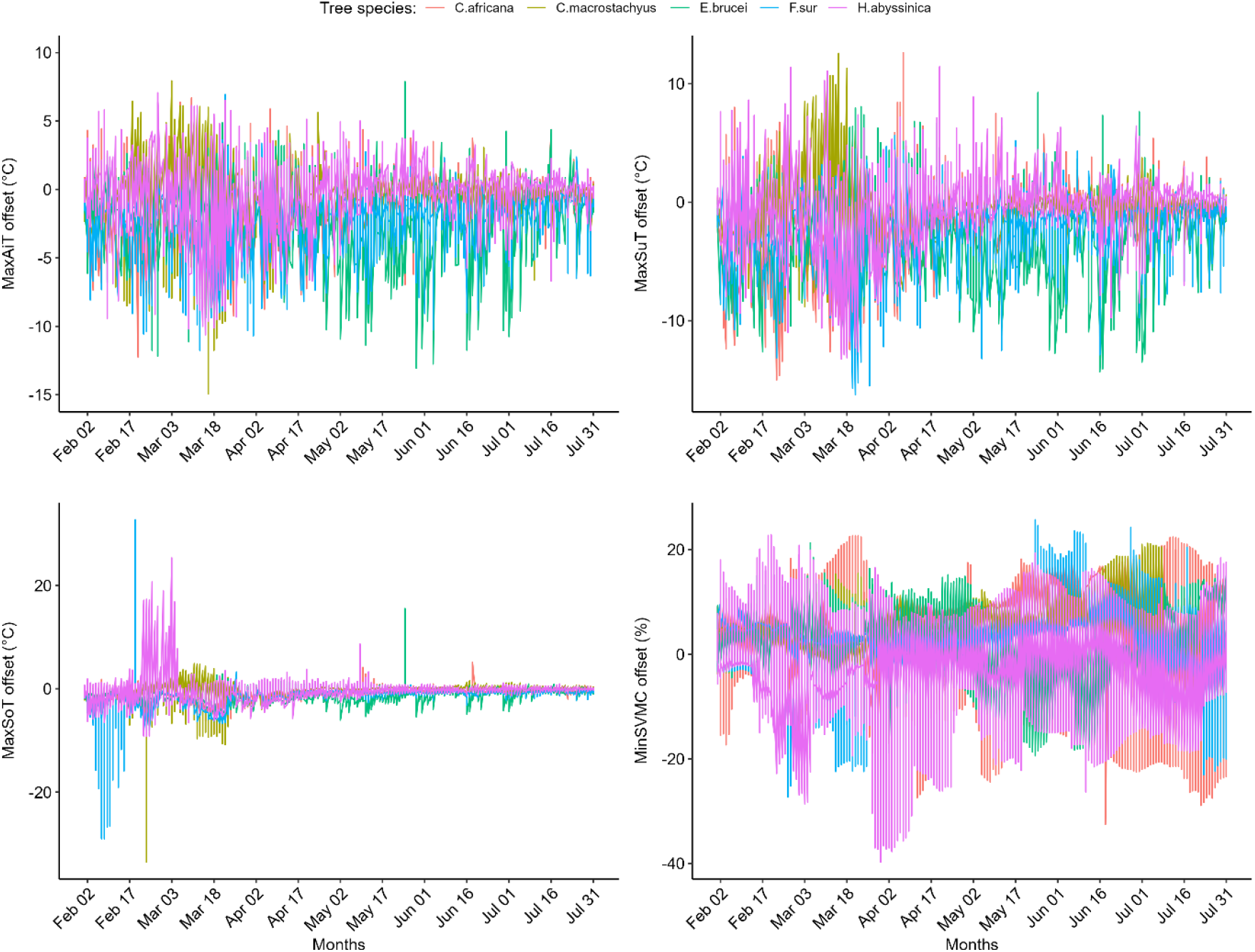
Microclimate offsets (maximum air, surface, and soil temperature, and soil volumetric moisture content) throughout the monitoring period (February 1, 2024, to July 31, 2024). AiT: Air temperature; SuT: Surface temperature; SoT: Soil temperature; and SVMC: Soil volumetric moisture content

